# Spatial scales of competition and a growth-motility tradeoff interact to determine bacterial coexistence

**DOI:** 10.1101/2022.01.05.474435

**Authors:** Thierry Kuhn, Marine Mamin, Saskia Bindschedler, Redouan Bshary, Aislinn Estoppey, Diego Gonzalez, Fabio Palmieri, Pilar Junier, Xiang-Yi Li Richter

## Abstract

The coexistence of competing species is a long-lasting puzzle in evolutionary ecology research. Despite abundant experimental evidence showing that the opportunity for coexistence decreases as niche overlap increases between species, bacterial species and strains competing for the same resources are commonly found across diverse spatially heterogeneous habitats. We thus hypothesized that the spatial scale of competition may play a key role in determining bacterial coexistence, and interact with other mechanisms that promote coexistence, including a growth-motility tradeoff. To test this hypothesis, we let two *Pseudomonas putida* strains compete at local and regional scales by inoculating them either in a mixed droplet or in separate droplets in the same Petri dish, respectively. We also created conditions that allow the bacterial strains to disperse across abiotic or fungal hyphae networks. We found that competition at the local scale led to competitive exclusion while regional competition promoted coexistence. When competing in the presence of dispersal networks, the growth-motility tradeoff promoted coexistence only when the strains were inoculated in separate droplets. Our results provide a mechanism by which existing laboratory data suggesting competitive exclusion at a local scale is reconciled with the widespread coexistence of competing bacterial strains in complex natural environments with dispersal.

## 2. Introduction

Bacteria are important components of the microbial community in diverse habitats, and the coexistence of different bacterial species and strains can serve important ecosystem functions, from soil nutrient cycling to providing defense against pathogens for their animal and plant hosts [1–5]. Competition experiments under controlled laboratory conditions revealed a general trend that the closer the phylogenetic distance between the competing species, the more likely that one species/strain competitively excludes the other, since niche overlap and antagonistic interactions are generally more intense between closely related species [6–8]. Nevertheless, competing species with overlapping niche use are commonly observed in natural habitats, including soils [9,10], fermented foods [11,12], and different parts of the human body [13–16]. Various mechanisms have been identified to promote the coexistence of bacteria [17], for example, metabolic niche partitioning [18–20], negative frequency-dependent selection [21–23], shared predators or parasites [24,25], and chemical warfare [26–28]. Many of these mechanisms involve tradeoffs between growth rate and other traits, such as the production and/or tolerance of antibiotics, resistance to adverse environments, and predator avoidance.

One understudied tradeoff is the tradeoff between bacterial growth and motility. This tradeoff can result from the metabolic costs of expressing and utilizing propulsion machinery (e.g., flagella) and genetic constraints that limit the simultaneous improvement of growth and motility during evolution [29–34]. The growth-motility tradeoff is interesting because it may help explain and reconcile the apparently conflicting results of a strong tendency of competitive exclusion found in laboratory experiments between competing species of overlapping resource niches and the widespread coexistence of intensely competing strains in nature. In contrast to the often spatially simple and well-mixed conditions in competition assays, natural environments such as the soil, skin, and fermented foods are featured by spatial heterogeneity that allows bacterial cells to interact at different spatial scales, and allow motile strains to disperse. Indeed, studies in animals and plants have shown that the competition-dispersal tradeoff (analogous to the growth-motility tradeoff in bacteria) can contribute to the coexistence of closely related species at relatively large regional spatial scales, but not at the smaller local scales [35,36]. Recent studies with bacteria also showed that a negative correlation between growth rate and dispersal ability can promote coexistence of closely related *Pseudomonas fluorescens* [37] strains, and *Escherichia coli* [32] strains. However, the experimental designs in these studies are highly simplified from biological reality. In the former study [37], the dispersal of bacteria was implemented by artificially transferring cell suspension across wells in 96-well plates at discrete time intervals; and in the later work [32], a simple and homogenous soft agar gel was used as the substrate where bacterial growth and dispersal took place. Therefore, there is need to test the effects of spatial scales of competition and the growth-motility tradeoff incorporating other relevant biological factors, for example, in spatially heterogeneous environments created naturally by fungal hyphae networks where bacterial cells have the opportunity to disperse by swimming by means of their flagella.

In this work, we study whether the spatial scale of competition can change the competition outcome between two closely-related *Pseudomonas putida* strains in the presence of a growth-motility tradeoff. We created heterogeneous environments by providing biotic and abiotic dispersal networks. The biotic networks were formed by living fungal mycelia [11,38,39], known as “fungal highways” in the soil environment [38]. The abiotic networks were based on glass fibers, which have been used in previous studies as an abiotic control for fungal hyphae [39–43]. In the absence of a dispersal network, the two bacterial strains compete either at the local scale when inoculated in mixed cell suspensions, or at the regional scale when inoculated in single-strain suspensions onto the same medium in a Petri dish. In the former case, direct contact between cells of different strains can occur, while in the latter case, cells of different strains do not touch each other but compete by utilizing a shared nutrient pool in the medium or can exchange volatile signals. In the presence of abiotic or biotic dispersal networks, the two bacterial strains compete at intermediate scales that were initially local/regional when inoculated in mixed/separate droplets, respectively. We performed experiments with a factorial design of three different spatial structures (i.e., no dispersal network, abiotic network, biotic network) and two initial scales of competition (i.e., local scale in mixed inoculation and regional scale in separate inoculation), and quantified the competition results in terms of whether the faster-growing strain can competitively exclude its slower-growing competitor.

## 3. Materials and Methods

### (a) Bacterial and fungal strains

For all experiments, we used two closely related flagellated strains of *Pseudomonas putida.* The KT2440 strain is a saprotrophic soil bacterium, originally isolated from the rhizosphere [44]; the UWC1 strain is documented as a spontaneous rifampicin-resistant mutant of the KT2440 [45]. A comparison between the genomes of the two strains showed additional differences, with a median identity of 95.8% between homologous genes (see Supplementary documents “genome_comparison.pdf” and “genome_comparison_table.xls”). The KT2440 and UWC1 strains were tagged with the green fluorescent protein (GFP) and mCherry, respectively, by Mini-Tn7 transposon insertions [46,47]. The different fluorescent labels enabled the two strains to be observed by epifluorescence stereomicroscopy and enumerated by flow cytometry. The fungal species used in the experiments was *Trichoderma rossicum* (NEU388) [43], a saprotrophic soil ascomycete that has been shown to allow various bacterial species (including *P. putida*) to move along its mycelial networks.

### (b) Culture conditions and inocula preparation

Bacterial cells were cryo-preserved in 30% (v/v) glycerol at −80 °C. To prepare the inocula, cells were first plated on Nutrient Agar (NA, 23 g/L, Carl Roth, AE92.2) supplemented with gentamycin (10 ppm) and grown at 30 °C in darkness. The strains were then sub-cultured once on NA through polygonal spreading and grown under the same conditions. Bacterial suspensions were prepared from overnight cultures grown in Nutrient Broth (NB, 25 g/L, Carl Roth, AE92.2) at ambient temperature under constant agitation at 120 rpm. Bacteria were collected from an overnight culture by centrifugation (3000 g for 10 minutes), washed once and resuspended in 0.01 M phosphate-buffered saline (PBS: 1.5 g/L Na_2_HPO_4_·2H_2_O, 0.25 g/L NaH_2_PO_4_·2H_2_O, 8,5 g/L NaCl, pH adjusted to 7.4). The cell densities of the two bacterial strains in the suspensions were determined by flow cytometry (see below) and then adjusted to the same value of 10^9^ cells/ml. After that, for the local competition scenario, a 1:1 ratio cell suspension was prepared by combining a proportion of the single-strain suspensions.

The fungus T. *rossicum* was maintained on a slanted agar tube at 4°C. It was plated on the malt-agar medium (MA: 12g/L malt extract Pulver amber, SIOS Homebrewing; 15g/L technical agar, BioLife Italiana) and incubated at 22 °C in darkness. Before being used for inoculation, the fungus was sub-cultured by replating a plug cored from the active apical margin of the two-day-old mycelial colony on a fresh medium and grown under the same conditions.

### (c) Experimental setup

As shown in Figure 1a, we inoculated the mCherry-labeled UWC1 strain and the GFP-labeled KT2440 strain in droplets of either single-strain or mixed cell suspensions onto plates of MA medium, under three different spatial conditions—no network, fungal mycelial network, and glass fiber network—each with four to eight replicates.

**Figure 1.**
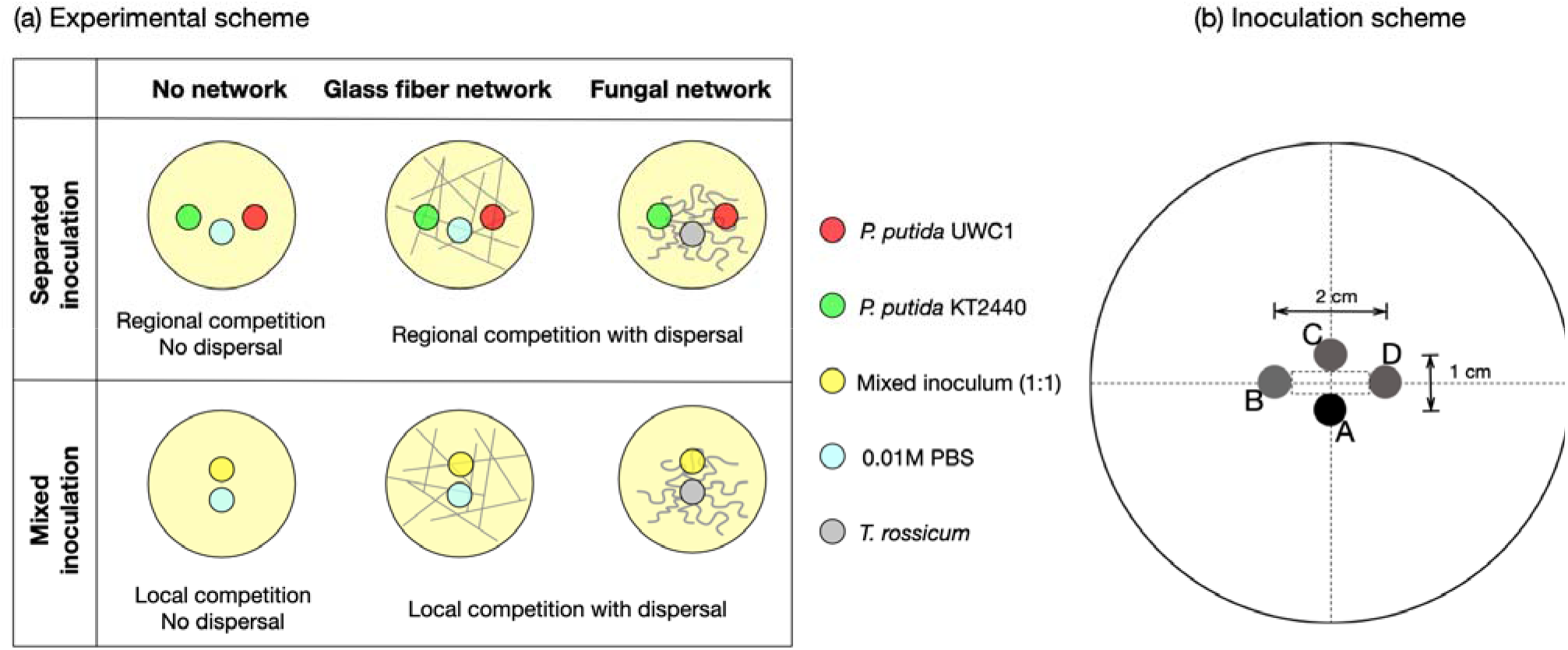
(a) Diagram of the experimental setup under spatial settings representing different spatial scales of competition. The inocula is represented in the diagram as disproportionally large for the purpose of a clear illustration. The diameters of the bacteria/PBS inocula and the fungal inocula were 5.8 mm and 5.0 mm, respectively. See panel (b) for a diagram of the actual positions of different inoculates on a □ 9 cm Petri dish. When applicable, the fugus or the PBS control were inoculated at position A, the KT2440 strain suspension was inoculated at position B, the mixed inoculum was inoculated at position C, and the UWC1 strain was inoculated at position D.

In the “no network” settings, both bacterial strains have limited capacity to disperse. The colonies (both separate or mixed) can only expand from the colony margin (Supplementary Figure S1). Under the “fungal network” and “glass fiber network” settings, the two bacterial strains used their flagella to swim and thereby dispersed in the liquid film along the surface of fungal hyphae or glass fibers, respectively [39].

In the network settings, when the two bacterial strains were inoculated separately, each strain could grow and start to disperse before encountering each other, and therefore the scale of interaction was initially regional. In this case, although the bacterial cells of the two strains were not in direct contact, they can still compete for resources diffused in the culturing medium in the same Petri dish. When inoculated in mixed droplets, the two strains engaged in direct competition right from the start before they could disperse, and thus the competition was initially local. Overall, the competition between the two strains occurred at finer scales when inoculated in mixed droplets, and the spatial scale of competition was coarser when the two strains were inoculated in separate droplets.

The bacteria and fungi inocula and the PBS control were positioned according to the diagram in Figure 1b. The distances between the inocula were determined based on preliminary tests. The relatively shorter distances between point A (where the fungi or PBS control were inoculated) and the other inocula ensures a quick start of bacterial-fungal interactions (the fungal mycelium network can grow past the bacteria inocula within 24 hours, see Supplementary Figure S2). The relatively larger distance between inocula B and D ensures that the separate colonies of the KT2440 and UWC1 strains did not get in direct contact by the end of the experiments (Supplementary Figure S3). We kept the positions of inoculation in Figure 1b fixed throughout all treatments to ensure that the fungal networks were of the same “age” when they expand and enter in contact with the bacterial colonies, because younger and older mycelia behave differently when interacting with bacterial colonies. Younger mycelial networks tend to stay outside bacterial colonies (Supplementary Figure S4a-b, fungal hyphae tips are in contact with bacterial colonies only at the colony margin) while older mycelial networks expand into the bacterial colonies (Supplementary Figures S4c-d, fungal hyphae tips are accessible to bacterial cells in the interior of the colonies). Because the UWC1 strain tends to constrain the KT2440 strain to the interior of a mixed colony (Supplementary Figure S1c), the KT2440 strain has a better chance to access and disperse on fungal networks (compare Supplementary Figure S5b and S5d, green cells were constrained to the interior of the bacterial colony in Figure S5b, but they were able to spread on the fungal hyphae in Figure S5d).

### (d) Inoculation procedures and culture conditions

In the “no network” setting, the inoculations of bacterial suspensions with separate/mixed strains and the PBS control were carried out by dropping on the medium surface 3 μl of the vortexed suspension with a microsyringe (Hamilton^®^ 1701N syringe 10ul needle size 26s ga) according to the corresponding positioning scheme in Figure 1. The resulting inocula were round-shaped with an area of 0.27 ± 0.011 cm^2^.

In the “glass fiber network” setting, we first placed around 30 glass fibers (8 μm diameter, 3-6 cm length, Cole-Parmer, USA) randomly on the agar plate and then performed the inoculations of bacteria as described earlier.

In the “fungal networks” setting, we first inoculated the fungus as a □ 5 mm plug cored with the wide-end of a Pasteur capillary pipette from the distal part of the T. *rossicum* mycelial network. Note that the mycelium was not in direct contact with the medium but laid on top of the plug. The two bacterial strains in separate or mixed droplets were inoculated immediately afterwards according to the schemes in Figure 1, following the same procedures as under the “no network”. To prevent the fungal inoculum agar plug from impeding the expansion of bacterial colony/colonies, we removed it after 24 hours, when the mycelial network had already expanded to the medium surface in the Petri dish.

After inoculation, the plates were parafilmed and incubated, in an upright position, at 22 °C in constant darkness.

### (e) Collection of bacterial cells for enumeration

After five days of growth, the bacteria cells were collected for enumeration by flow cytometry. Under the “no network” settings, we retrieved all bacterial cells by scraping the whole colony and resuspending the cells in 0.22 μm-filtered PBS. The suspensions were then stored on ice.

Under the “glass fiber network” and the “fungal network” settings, we first removed the bacterial colonies from the plates with a scalpel. We excluded cells in the bacterial colonies because we aimed to examine the relative abundance of the two bacterial strains that had already dispersed in the fungal/glass fiber networks. To collect those cells, we added 7 ml filtered PBS to the Petri dish, closed it, and agitated it at 100 rpm for 20 minutes on a rotary shaker. After that, we intensively flushed the surface of the medium with a 1000 pl micropipette, collected 5 ml of the resulting suspension, and stored it on ice.

All the suspensions were then sonicated in a sonic bath at 35 kHz (Transsonic 310, Elma) twice for 30 seconds each and with a 30-seconds break in between [38]. This step was done to disaggregate bacterial clumps and/or to separate cells that might have stuck to the fungal/glass fiber fragments that might present in the suspension by chance. To remove those fragments before the flow cytometry analysis, we filtered the suspension through a polycarbonate filter with pores of 10 μm (Whatman, Nucleopore Track-Etch Membrane) pre-autoclaved in the Swinnex filters holders. All samples were kept at 4 °C before performing the flow cytometry analysis the day after.

### (f) Flow cytometry

We estimated the relative abundances of the bacterial strains with a CyFlow space cytometer (Partec-System) equipped with a blue laser (480 nm/20mV) and a green laser (532 nm/30mV). The FSC and SSC optical parameters are associated with the blue laser. The green fluorescence of the KT2440 strain was excited by the blue laser and detected on a photomultiplier tube with a 488-nm bandpass filter (FL1 parameter). The red fluorescence of the UWC1 strain was excited by the green laser and detected on a photomultiplier tube associated with a 590/50-nm bandpass filter (FL2 parameter). The lasers were arranged in parallel, with a delay of 50 μs between the two signals. Calibration of the instruments was verified before the analysis with calibration beads (Partec-Sysmex). The leading trigger was set on SSC, with a lower threshold of 500. Gains were set to 700 for FL1 and 750 for FL2. A four-decade logarithmic amplification was used for all parameters. The instrument was equipped with a true volumetric absolute counting facility: events were counted in a defined volume of 200 μl. A speed of 2 μl/s was used. We diluted the samples with 0.22 μm-filtered PBS to reach an event rate of ~1500 total events/s. This rate was determined with preliminary assays to yield an optimal signal to noise ratio. Samples were kept on ice during the whole analysis and were run in random order. Analysis of the flow cytometry data was performed with the instrument software FloMax. Events corresponding to the mCherry and GFP fluorescent signals were selected and counted with a quadrant gate set on a biplot of the FL1 and FL2 parameters.

### (g) Data preparation and statistics

We focused on the relative abundance of the UWC1 and KT2440 strains, and therefore, we calculated the cell density ratio in each replicate at the end of the experiments. We then transformed the data by taking the common log (of base 10) and confirmed the normality of the transformed data with the Shapiro-Wilk normality test. We then performed the Student’s *t*-test against our null hypothesis that the relative abundance of the two strains remains identical to that at the beginning of experiments (with a log ratio of zero). The competitions between the abundances of the two strains when inoculated in mixed or separate droplets were performed by the Welch’s two sample *t*-tests. The statistical significance threshold value was chosen at p = 0.05. We adjusted the p-values to account for the false discovery rate due to multiple comparisons using the Benjamini and Hochberg method [48]. All statistical analyses were carried out within the R statistical environment [49]. The R scripts are provided in the Supplementary Information.

## 4. Results

Under the “no network” treatment, when bacteria cells were inoculated in separate droplets (corresponding to regional competition), the relative abundance of the KT2440 strain and the UWC1 strain remained similar (*p* = 0.569) after 5 days of growth (Figure 2a, Tables 1 and 2). In contrast, in mixed inocula (corresponding to local competition) the UWC1 strain reached much higher cell proportions (*p* < 0.01) at the end of the experiments (the UWC1 strain was on average 5.43 times more abundant than the KT2440 strain), showing a growth advantage of the UWC1 strain in direct competition. Moreover, the spatial distribution of the two fluorescently tagged strains indicated that the UWC1 strain has expanded faster and constrained the KT2440 strain to the interior of the mixed colony, thus blocking the further growth of the KT2440 strain (Figure 2b). The marked difference in the relative competitiveness of the two strains implies the competitive exclusion of the KT2440 strain by the UWC1 strain at the local scale in the long term.

**Figure 2.**
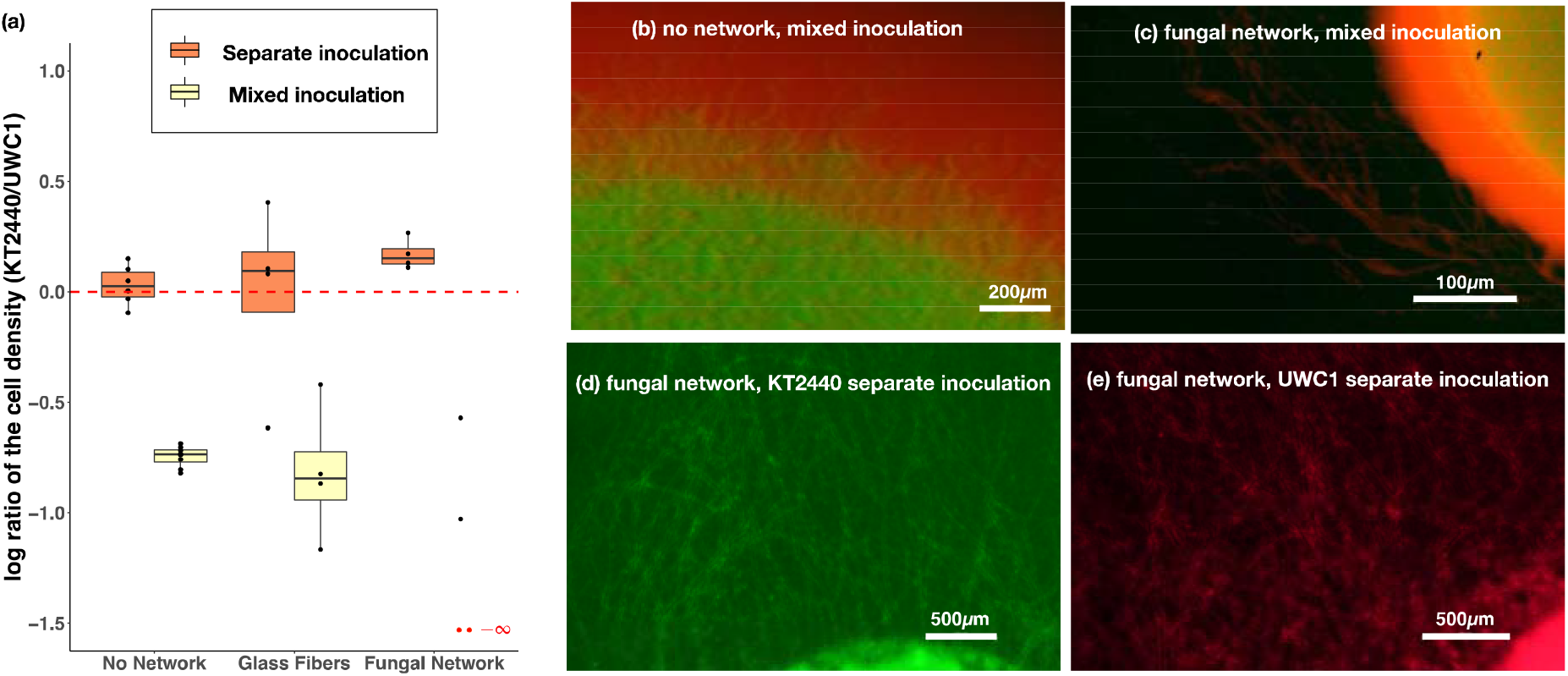
Box-Whisker plots representing the distribution of the common logarithm (base 10) of the cell density ratio (KT2440/UWC1) under different experimental settings. The KT2440/UWC1 strain is more abundant than the other strain above/below the red horizontal line at log ratio of 0, respectively. The boxplot center lines show the median, box limits show upper and lower quartiles, whiskers show 1.5x interquartile range, and each point corresponds to the result of an independent replicate. The red dashed line represents equal cell densities of the two bacterial strains. The numbers of replicates were 6 and 8 under the “no network” setting for the separate and mixed inoculations, respectively. Under all other settings, the numbers of replicates were 4. Competitive exclusion of the KT2440 strain by the UWC1 strain occurred in two replicates under the fungal network treatment with mixed strain inoculates (the log density ratio was —∞, marked with the two red dots). (b) Colony formed by bacterial strains in a mixed droplet two days after inoculation in the absence of a dispersal network. (c) Colony formed by bacterial strains in a mixed droplet in the presence of fungal hyphae networks two days after inoculation. Only the UWC1 strain with red fluorescence was visible along the fungal hyphae. (d) Dispersed KT2440 cells along fungal hyphae networks three days after being inoculated in a separate droplet. (e) Dispersed UWC1 cells along fungal hyphae networks two days after being inoculated in a separate droplet. The pictures were taken at magnifications of 10x, 3x, 5x, and 5x with epifluorescence stereomicroscopy in panels b, c, d and e, respectively.

**Table 1.** The median of the absolute density ratio (before log transformation) of the KT2440 strain relative to the UWC1 strain.

|  | No network | Glass fiber network | Fungal network |
| --- | --- | --- | --- |
| Separate inoculation | 1.064 | 1.239 | 1.418 |
| Mixed inoculation | 0.184 | 0.143 | N.A.<sup>1</sup> |

<sup>1</sup> The KT2440 strain was competitively excluded in two of the four replicates.

**Table 2.**
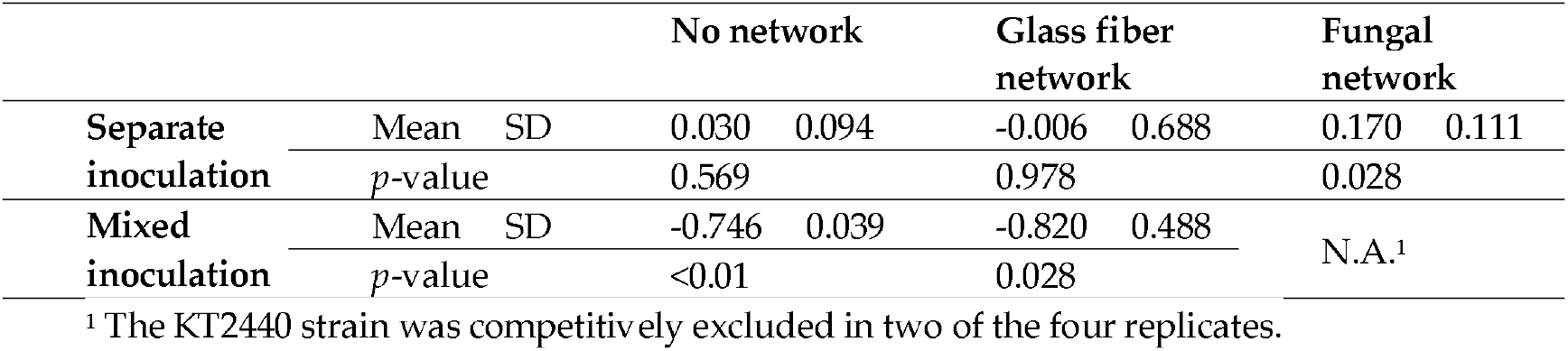
Mean and the standard deviation of the log density ratio between the KT2440 and the UWC1 strain, and the p-value of the *t*-test with a null hypothesis that the mean of the log-ratio was not significantly different from zero.

Under the “glass fibers” treatment, the KT2440 strain was slightly more abundant (but the difference was not statistically significant, *p* = 0.978) when the two bacterial strains were inoculated separately (corresponding to an intermediate scale of competition that was initially regional, see Figure 2a, Tables 1), reaching 1.239 times the density of the UWC1 strain. We therefore cannot reject the null hypothesis that the relative abundance of the two strains remained identical (Table 2). This result implies possible long-term coexistence of the two strains at the regional scale. In plates where the two bacterial strains were inoculated in mixed droplets (corresponding to an intermediate scale of competition that was initially local), similar to the results of local competition under the “no network” treatment, the UWC1 strain has a substantial competitive advantage (*p* = 0.028) and reached on average 6.99 times the density of the KT2440 strain, implying that competitive exclusion was likely at the local scale (Figure 2a, Tables 1 and 2).

Under the “fungal network” treatment, the KT2440 strain reached even larger (statistically significant, *p* = 0.028) relative abundances when inoculated in separate droplets (1.42 times the abundance of the UWC1 strain), implying a competitive advantage at the regional scale. The competitive advantage of the KT2440 strain outside the initial inocula resulted from its higher motility and faster spread in the presence of fungal hyphae networks. See Supplementary Figure S7 for direct verification of the motility advantage of the KT2440 strain. The distribution of bacterial cells in the fungal network can be considered as mini-colonies with either mixed or separate strains. Since the KT2440 strain grows equally fast as the UWC1 strain in separate colonies, and it grows much slower than the UWC1 strain in mixed colonies, the overall higher densities of the KT2440 strain was likely to result from its superior motility in the presence of dispersal networks provided by fungal hyphae. In mixed inoculations, however, the KT2440 strain was competitively excluded outside the initial inocula (Figure 2c for an example) in two of the replicates (in these cases, the cell densities of the KT2440 strain were so low that they were not distinguishable from the noise background in the flow cytometry assays), and in the other two replicates, the cell densities of the UWC1 strain was 3.72 times and 10.67 times more abundant than the KT2440 strain (Figure 2a, Tables 1 and 2). The spatial distribution of the two tagged strains indicated that when inoculated in mixed droplets in the presence of fungal networks, the slower-growing KT2440 strain was constrained to the interior of the mixed colony, and consequently, was not able to disperse through the fungal networks (Figure 2c, note that only red fluorescence was visible along the fungal hyphae connected to the colony). In contrast, when inoculated in separate droplets, both bacterial strains can efficiently disperse along fungal hyphae networks (Figure 2d-e).

Across all spatial settings, the relative abundance of UWC1 and KT2440 strains differ greatly depending on whether they were inoculated in separate or mixed droplets (Figure 2a, Table 3), showing the importance of the spatial scale of competition in determining the competition outcome and long-term coexistence of the two bacterial strains, in the presence or absence of dispersal networks.

**Table 3.** Comparison between the distributions of the common logarithm (base 10) of the cell density ratio (KT2440/UWC1) under separate and mixed inoculation treatments.

|  |  |
| --- | --- |
| Spatial settings | p-value of the Welch's two sample t-test |
| No network | <0.001 |
| Glass fiber network | 0.025 |
| Fungal hyphae network | N.A., because of the two values under mixed inoculation treatment |

## 5. Discussion

Our experimental results showed a clear pattern: local competition between the two bacterial strains (treatments with mixed inocula) caused competitive exclusion, while competition at the regional scale (treatments with separate inocula) is more favorable for coexistence. This is consistent with the effect of the spatial scale on the competition outcome found in larger organisms [35,36]. Our results thus help to reconcile the conflicting findings that bacteria [6–8] and protists [50] species/strains that overlap in their niches of resource utilization tend to competitively exclude each other in well-mixed liquid cultures (implying local competition), and the widespread phenomenon of microbial species/strains coexisting while competing for the same resources in spatially complex environment (implying regional competition), such as cheese rind [11,12], the human gut [13,16] and skin [14]. It is important to note that spatial complexity does not automatically promote biodiversity. The effect of spatial structure crucially depends on the type of interactions between species, and in some cases, adding spatial structure can lead to the breakdown of diversity and competitive exclusion of species that could have coexisted under a well-mixed environment [51–53].

In the current work, we showed that when the competing species are involved in a growth-motility tradeoff, spatial structure provided by dispersal networks can help the competing species to coexist. In the presence of dispersal networks, the higher motility of the slower-growing strain (KT2440) can confer an advantage only when the initial scale of competition was regional (the two strains were inoculated in separate droplets). Otherwise, the faster-growing strain (UWC1) can quickly outgrow the slower-growing strain, constraining the latter to the interior of the mixed colony, and thus preventing it from dispersing. Instead of comparing the growth rates of the UWC1 and KT2440 strains in isolation, we inferred the growth advantage of the UWC1 strain in direct competition by comparing the log ratios of the cell densities under mixed and separate inoculations in the “no network” setting, controlling that spatial arrangement was the only difference. Microbial growth rates are highly context-dependent. Literature has shown that even when growing in monoculture the relative growth rates of competing species can switch in different environments (e.g. *in vivo* in the host environment and *in vitro* when cultured in microcosms [54]), and the relative growth rates in coculture can show complex non-linear patterns with multiple crossings at different initial frequency combinations [22]. Not only the growth rate, the motility of microorganisms is also context-dependent and can be sensitive to environmental conditions [55]. Because of the strong association between microbial growth and dispersal rates and their sensitivity to biotic and abiotic environmental factors, it is important to measure them in the same competition context, as we did in the current work.

Note that the significantly higher abundance of the KT2440 strain in the presence of fungal networks when inoculated in separate droplets does not imply that it can competitively exclude the UWC1 strain for three reasons. First, due to the limitations of the experimental system (e.g., the fungal network has already covered the entire Petri dish and could not expand further by the end of five days), our results only showed a snapshot of the competition dynamics and therefore have limited power to predict the entire evolutionary trajectory. In addition, we focused on quantifying the relative abundance of cells *outside* the colonies formed by the initial inocula, and thus the result of a higher abundance of the KT2440 cells was limited to cells descended from the dispersed ones, and it is possible that the UWC1 strain has nevertheless reached higher abundance *inside* the initial colony. Finally, our results have shown a consistent growth advantage of the UWC1 strain when competition occurs locally. Therefore, we expect that as soon as the cells of the two strains meet in the dispersal network and engage in direct local competition, the UWC1 strain would overgrow its competitor locally. The competitive advantage of the UWC1 strain in local competition is highly contingent on its ability to block the KT2440 strain from accessing the dispersal network. Unlike diffusible nutrients, space is a non-sharable resource. By occupying specific locations in space, such as the expansion frontiers of a mixed bacterial colony or the locations where a mixed colony meets a dispersal network, the faster-growing species (the UWC1 strain in our case) not only prevents its competitor from occupying the same space per se, but also deprives it the opportunity to use a section of space extending from the colony margin [56–58], and in the presence of a dispersal network, also the section of space in the network that is accessible through that location. Existing work has shown that physical interactions play essential roles in shaping the structure of microbial colonies and biofilms [59–61]. In this work, we showed that the spatial occupation of the UWC1 cells at the “entrance points” of a dispersal network gives them an “opportunity advantage” to explore and expand to new territories.

The competitive advantage of the KT2440 strain when dispersal is allowed suggests that the growth-motility tradeoff plays a role in promoting the coexistence of the two bacterial strains, with the UWC1 strain growing faster under direct competition while the KT2440 is better at dispersing and exploring empty habitats. When competing species are regulated by a tradeoff between their rate of reproduction and their ability to disperse, theory predicts that there are conditions where coexistence is possible [62,63]. A previous study with two *P. fluorescens* strains also demonstrated that regional competition in the presence of a growth-dispersal tradeoff can promote coexistence [37]. In the aforementioned study, however, dispersal occurred by manually transferring bacterial cells at discrete intervals. Such experimental design, while promoting the controllability of the system and helping to reduce variation between different replicates, is highly artificial and corresponds to natural conditions that only rarely occur (e.g., occasional heavy rains that saturate the soil for a short time [47,64,65]). In this work, we allowed continuous dispersal of the two bacterial strains, either through glass fiber networks, or fungal mycelial networks that can interact dynamically with the bacteria [66,67]. The bacterial dispersal in our experiments thus corresponds to that in environments such as the typical well-drained soil, where the movement of bacteria is restricted by the thinness and patchiness of the liquid films [68] on the surface of abiotic (e.g., rocks and soil particles) and biotic (e.g., roots, fungal hyphae) components. Despite the additional factors and variation involved, our results showed robustly that interactions at the local scale tend to cause competitive exclusion, while coexistence is more likely when competition is regional.

In this work, the abiotic network was formed by placing thin glass fibers randomly on a Petri dish, while the biotic network was formed by the actively growing fungal hyphae, which follow species-specific structural patterns. The glass fiber network was designed as an abiotic “control” of the fungal network as in several previous studies [39–43]. However, our work showed that the glass fiber network is not an ideal abiotic substitute for the fungal network, partly because the liquid films formed around glass fibers are much thicker than those surrounding fungal hyphae (Supplementary Figure S6). The thickness of liquid film determines whether cells can dispersal solely actively by flagella-propelled swimming or also passively driven by hydraulic flow in the liquid film [69]. The hydraulic flow caused by inoculation may have flushed some KT2440 cells into the dispersal network, leading to a large variation in the competition outcome under the glass fiber treatment. Another factor contributing to the variation could be the randomness of dispersal network created when placing the extremely thin (8 *μm* in diameter) and brittle glass fibers on the Petri dish (Supplementary Figure S6). Theory and experiments have shown that the network structure and connectivity can shape biodiversity patterns [70,71]. Since we were not able to precisely control the abiotic network structures in the current work, large variations persist in the relative abundance of the two bacterial strains. This also limits our ability to investigate deeper the interactions between network type and different ways of inoculation, for example, by performing a two-factor ANOVA test. Therefore, a valuable next step would be to tease apart the large variations of our current results under the abiotic and biotic network settings by manipulating their structures, for example, by using 3D printed abiotic hyphae models with fixed structure and surface physicochemical properties, or microfluidic platforms that constrain the growth of fungal hyphae networks into pre-defined topology [72,73].

## 6. Conclusion

To conclude, using experiments with two closely related strains of the bacteria *Pseudomonas putida,* we found that spatial scales of competition and the growth-motility tradeoff interact to determine the competition outcome. We showed that local competition caused competitive exclusion while regional competition is more favorable for coexistence. The results are consistent with previous findings in animals and plants. Furthermore, we showed that in the presence of dispersal networks, the growth-motility tradeoff can promote coexistence only when the scale of competition was regional initially. Only in this case, the slower growing but faster dispersing strain has the opportunity to use the dispersal networks to escape from direct local competition with its faster-growing competitor. It is interesting to ask whether the phenomenon we observed in this study may generalize to microbial competition dynamics in other spatially complex environments, where interactions occur at different spatial scales and in the presence of dispersal networks provided by diverse fungi and fungi-like organisms. Therefore, we encourage future work to systematically study the interactions between spatial scales of competition and the growth-motility tradeoff between pairs of microorganisms with different phylogenetic relatedness. The current work identified new factors that affect microbial competition dynamics in spatially complex environments — in particular, along dispersal networks provided by fungal hyphae, and raised new questions regarding the roles of network topology and the biotic interactions between fungi and different bacterial strains that disperse along them. Future investigations along these lines would help us delve deeper into the mechanisms that govern microbial competition dynamics and coexistence in spatially complex environments.

## Supporting information

Appendix containing supplementary figures and tables

## Acknowledgments

The authors would like to thank Arnaud Dechesne and Jan van der Meer for providing the two bacteria strains.

## Ethical Statement

Not applicable.

## Data Accessibility

All data supporting the reported results can be found in the data tables in the manuscript and the Supplementary Materials.

## Authors’ Contributions

P.J., R.B, S.B. conceived the research project, P.J., R.B., X.L., S.B., D.G. conceptualized the experiments, M.M. and T.K. developed the methodology, performed the experiments and analyzed the data, A.E. and F.P. provided supports to the methodology, A.E. and D.G. helped with the flow cytometry assays, X.L. wrote the manuscript with inputs from all authors.

