## Appendix containing supplementary figures and tables for "Spatial scales of competition and a growth-motility tradeoff interact to determine bacterial coexistence"

### Supplementary Materials

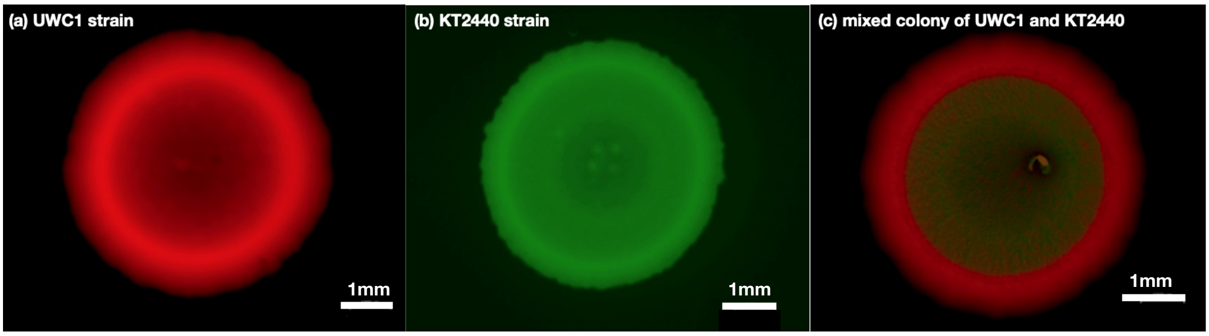


**Figure S1.** The UWC1 and KT2440 strains in separate or mixed colonies in the absence of a dispersal network. The pictures were taken with an epifluorescence microscope 2 days after inoculation.

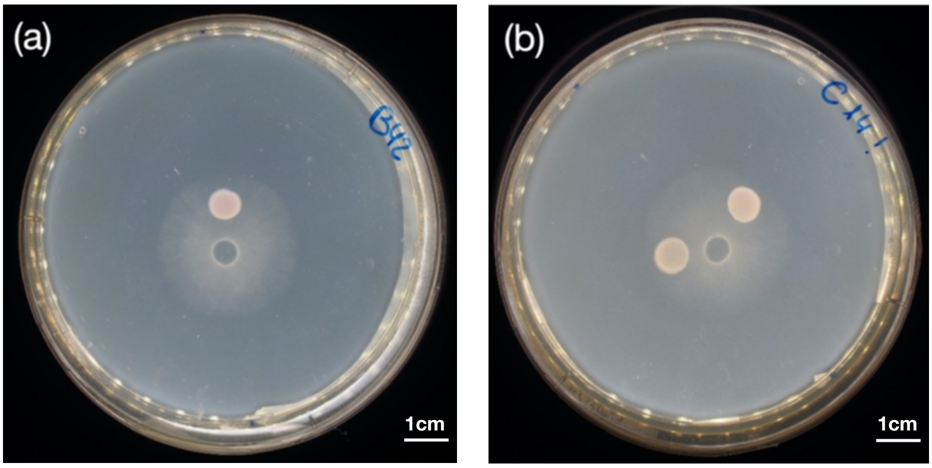


**Figure S2.** Two examples of Petri dishes with bacteria and the fungus *Trichoderma rossicum* showing that the latter has already grown passed the bacterial inocula after 24h of growth, either in the case of a single bacterial inoculum (panel a, bacterial strain was UWC1) or two separated inocula (panel b, bacterial strains were UWC1 and KT2440). The pictures were taken after having removed the fungal inoculum plug.

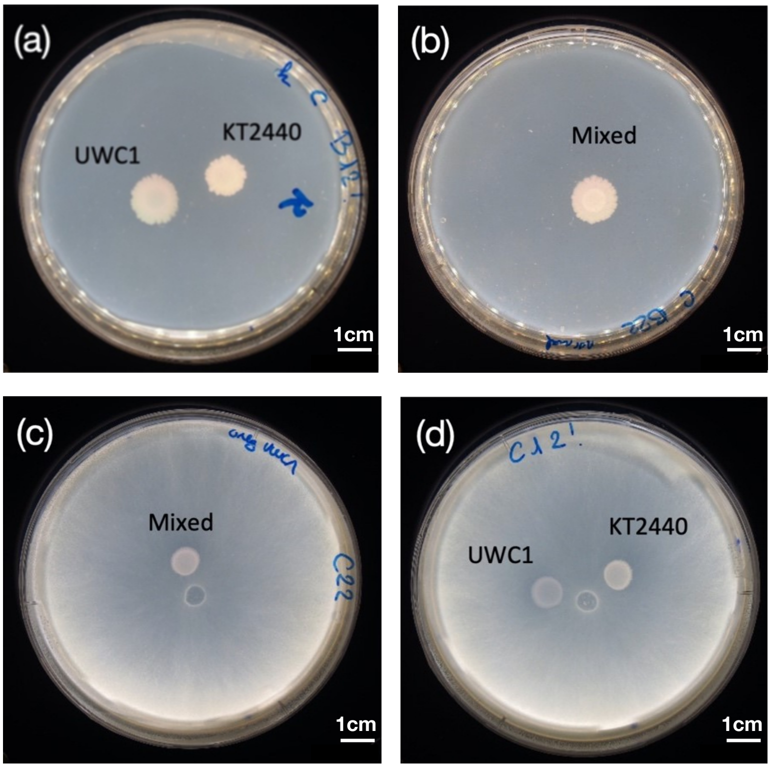


**Figure S3.** Examples of Petri dishes after 5 days of growth showing that the margins of the bacterial colonies remain separate at the end of the experiments. Dispersal networks formed by the fungus *Trichoderma rossicum* were absent in panels (a) and (b), and present in panels (c) and (d).

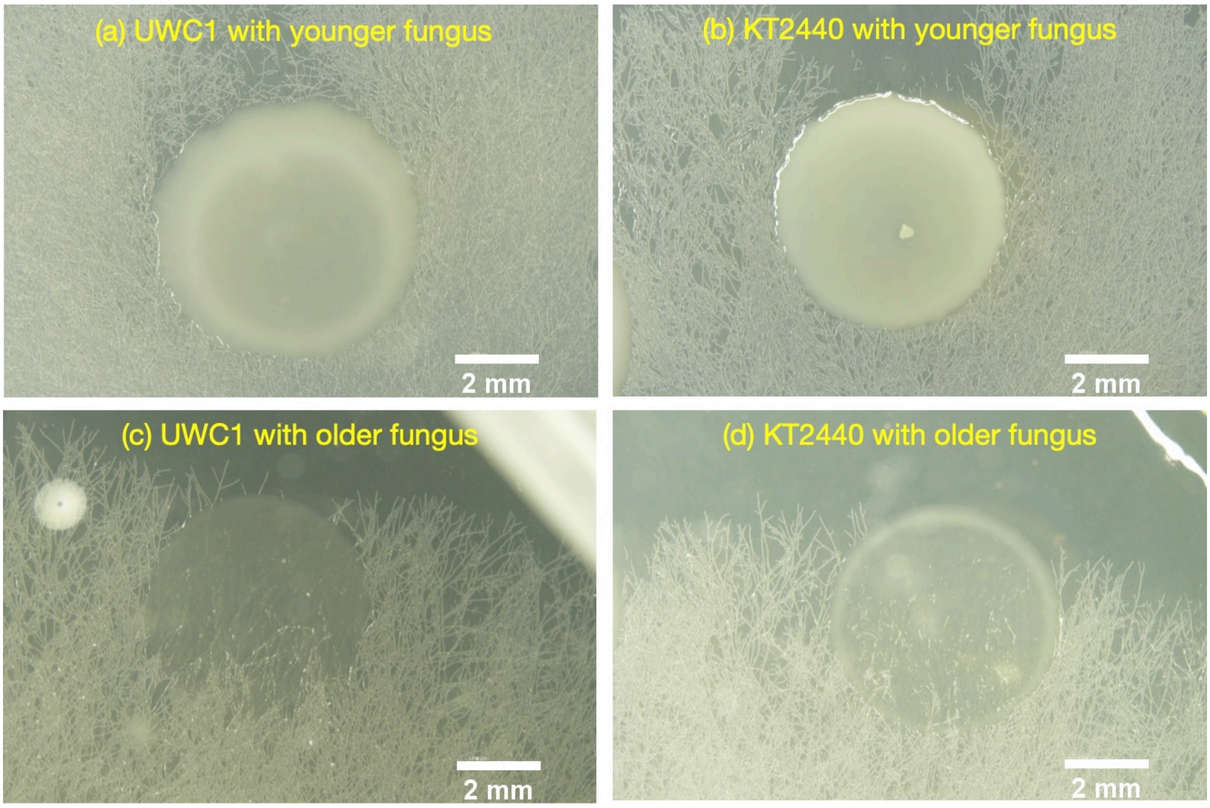


**Figure S4.** Bright field photos of the colonies of the UWC1 and KT2440 strains in the presence of younger (2 days) and older (5 days) fungal networks. The younger fungus does not invade the bacterial colonies, while the older fungus penetrated the colonies of both bacterial strains.

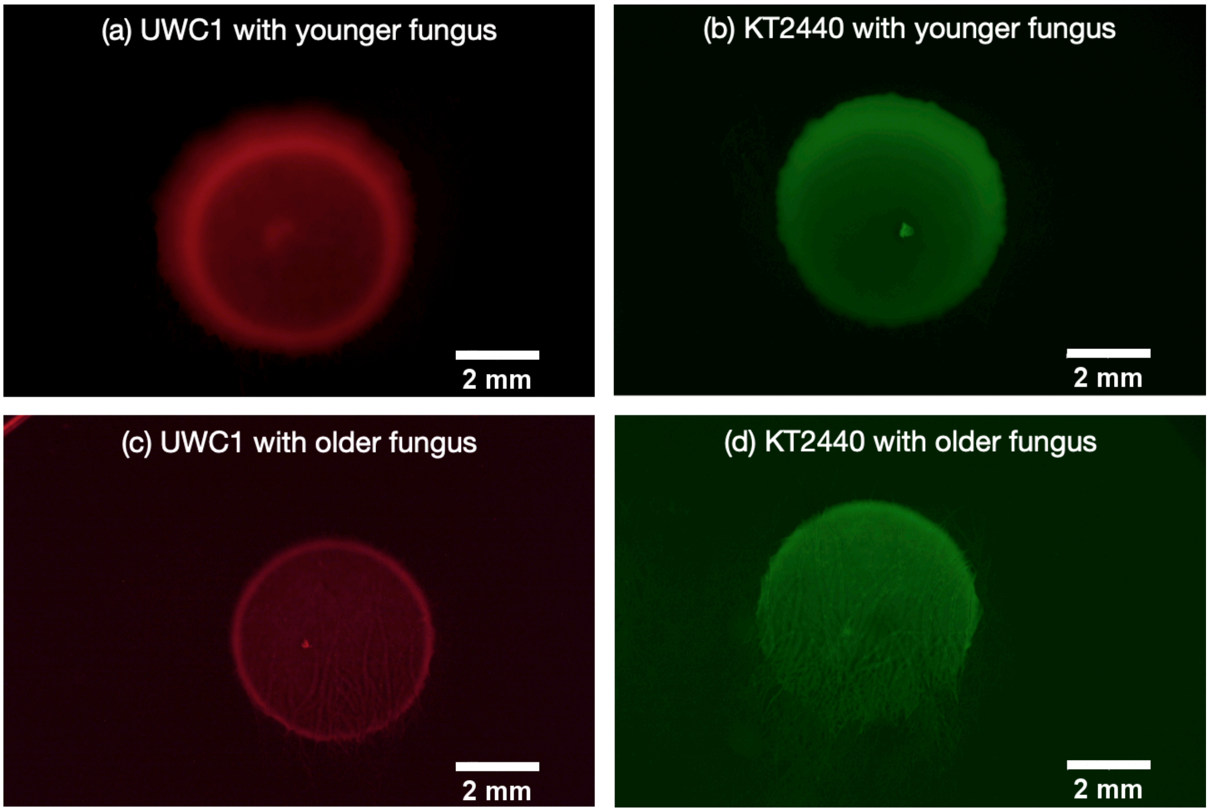


**Figure S5.** Pictures of the same plates as in Figure S3 taken with epifluorescence microscopy, showing the spatial distribution of bacterial cells along fungal hyphae networks. The UWC1 strain can spread efficiently on younger fungal networks, without the fungal hyphae penetrating the colony (panel a), but it spread less well on older fungal networks (panel c), despite that the fungal network has penetrated the colony. The effects of younger and older fungus were the opposite for the KT2440 strain – bacterial cells did not spread on the networks of younger fungus (panel b), but dispersed efficiently along the networks of older fungus that penetrated the bacterial colony (panel d).

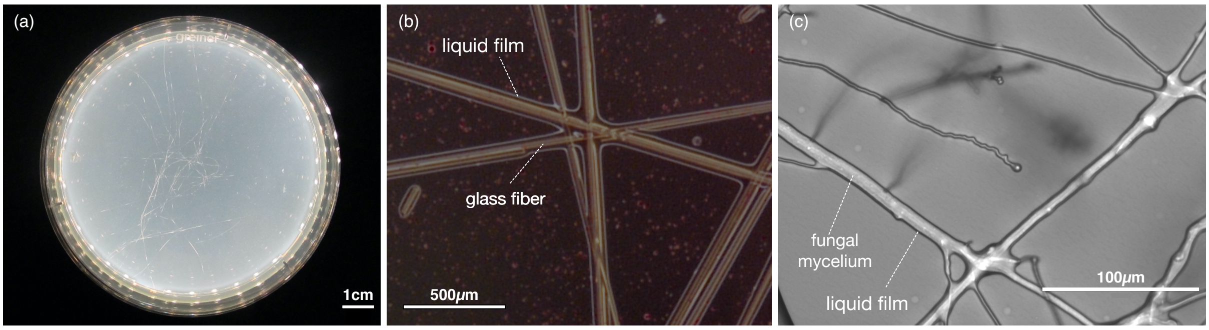


**Figure S6.** (a) Picture of a glass fiber network placed on the culturing medium in a Petri dish. Around 30 pieces of fine (8 $\mu m$ diameter) glass fibers were distributed randomly to form the network. (b) Relatively thick liquid films form along the surface of glass fibers. (c) Thin liquid films form along the surface of fungal mycelia.

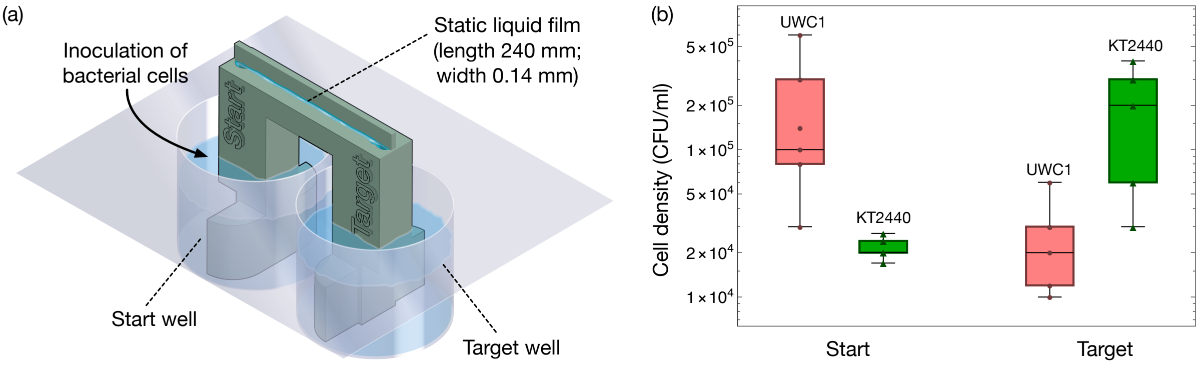


**Figure S7.** (a) Illustration of the design of bacterial motility assay. 5 $\mu l$ cell suspension of the UWC1 or KT2440 strain (adjusted to an optical density of one) were added to the start well of the "bacterial bridge" device (described in detail in Kuhn *et al*, 2022), where two medium reservoirs were connected by a static liquid film through two vertical capillaries. The culture medium contained 0.01 M PBS with 1% of NB and 1% of Ficoll (10g/L, Sigma-Aldrich). Due to a lack of hydraulic flow in the liquid film, bacterial cells can move from the start well to the target well only by active flagella-propelled swimming. Cell densities in the start and target wells were measured 72 hours after inoculation. Experiments for each strain have six independent replicates. (b) Box-Whisker charts showing the cell densities in the start and target wells at the end of experiments. More cells of the KT2440 strain dispersed from the start well to the target well than cells of the UWC1 strain ($p$ = 0.025, Student $t$-test).

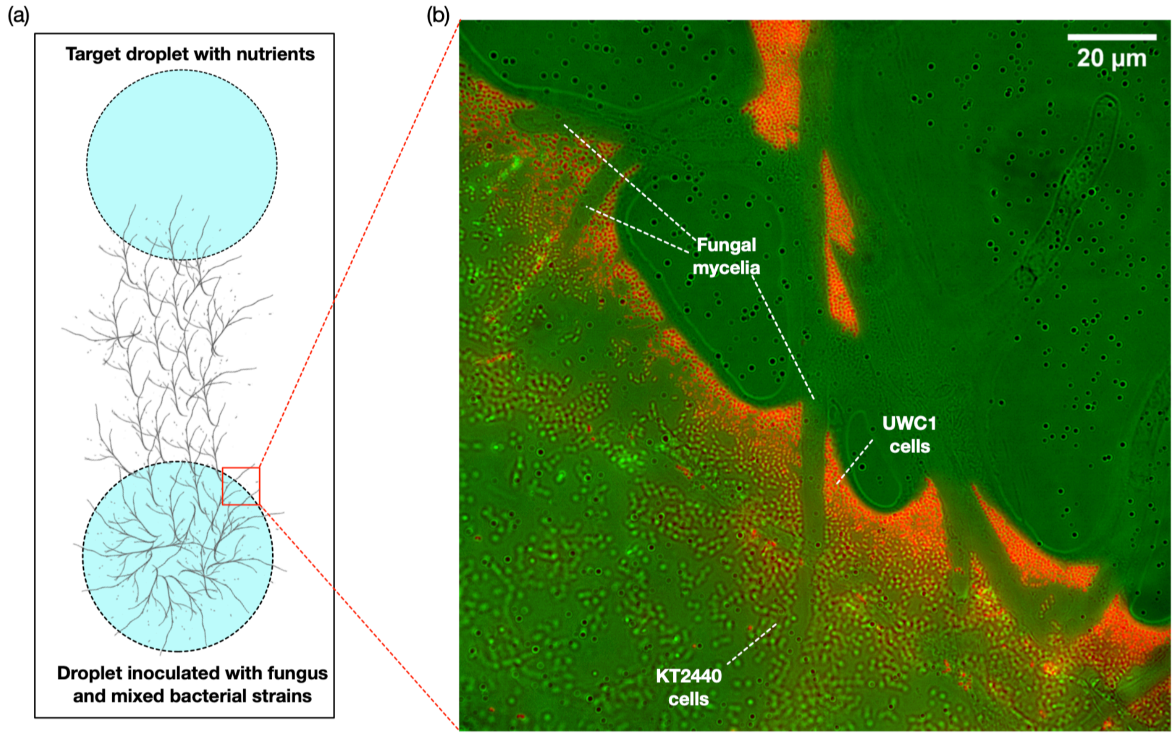


**Figure S8.** (a) Illustration of bacterial dispersal in the fungal network on a glass slide. The oomycete *Pythium ultimum* and a 1:1 mixture of *Pseudomonas putida* KT2440 and UWC1 cells were inoculated in a droplet of NB medium. Within 72 hours, the hyphal network has reached a separate target droplet. (b)Epifluorescence microscopic picture showing the advantage of the UWC1 strain (red cells) in accessing the fungal mycelial network on the glass slide. The upper left — lower right diagonal of the figure corresponds to the margin of the source droplet where the fungus and bacterial strains were inoculated. The picture was taken 72 hours after inoculation.

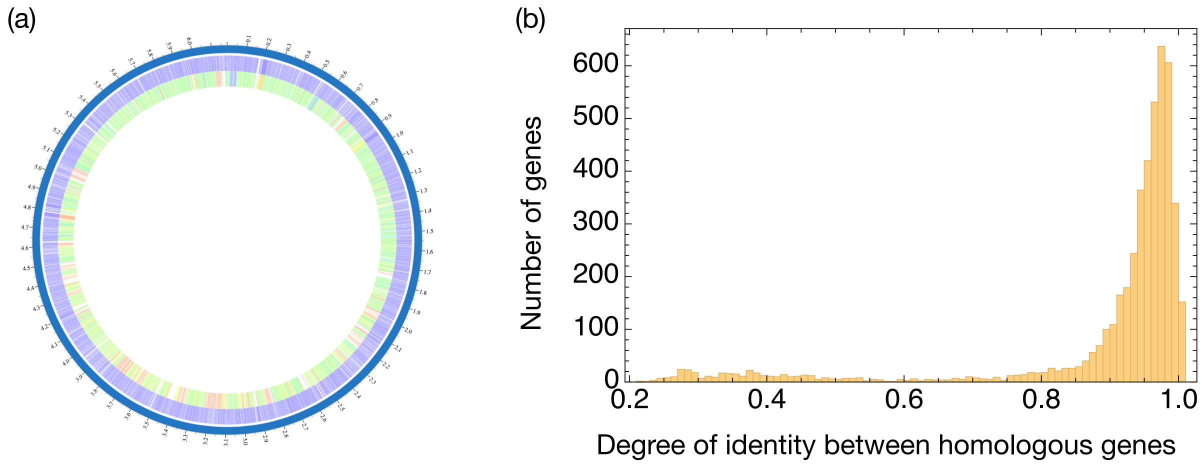


**Figure S9.** (a) A snapshot of the genome comparison between the KT2440 and UWC1 strains. Annotated reference genome *P. putida* KT2440 (Taxonomy ID: 160488) was compared to annotated representative genome *P. putida* NBRC 14164 (synonymous to strain UWC1; Taxonomy ID: 1407054) using the proteome comparison tool provided by (https://patricbrc.org; Davis et al, 2020). Both genomes have annotated versions already available on the NCBI genome database website. The color scale from purple to red represents genome identity from high to low. The tracks from outside to inside are the KT2440 and UWC1 strains, respectively. See supplementary file "genome_comparison.pdf" and supplementary table "genome_comparison_table.xls" for detailed information. (b) Histogram of the identity levels between homologous genes. The chart represents a total of 4691 genes.
